# Blocking the Necroptosis Pathway Decreases RPE and Photoreceptor Damage Induced by NaIO_3_

**DOI:** 10.1101/387068

**Authors:** Haijiang Lin, Miin Roh, Hidetaka Matsumoto, Alp Atik, Peggy Bouzika, Albert Alhatem, Joan W. Miller, Demetrios G. Vavvas

**Author notes:** These authors contributed equally to the work presented here and should therefore be regarded as equivalent authors (Haijiang Lin, Miin Roh). Corresponding author: Demetrios Vavvas. Department of Ophthalmology, Department of Ophthalmology, Massachusetts Eye and Ear Infirmary, Harvard Medical School. 243 Charles Street, Boston, MA 02114.

## Abstract

**Purpose:** Sodium iodate (NaIO3) has been extensively used as a retinotoxin to induce RPE cell damage and degeneration of photoreceptors *in vitro* and *in vivo*. RIP-Kinase dependent programmed necrosis is an important redundant cell death pathway involved in photoreceptor cell death. We wanted to determine whether these pathways are actively involved in RPE and photoreceptor cell death after NaIO3 insult.

**Methods:** ARPE-19 cells were exposed to different concentrations of NaIO3 in the presence or absence of various concentrations of a RIPK inhibitor (Nec-1) or a pan-caspase inhibitor (Z-VAD), individually or combined. Cell death was determined at different time points by MTT (Sigma-Aldrich), LDH (Promega) and TUNEL (Millipore) assay. C57BL/6 and RIP3^−/-^ mice were treated with a peritoneal injection of NaIO3 and eyes were enucleated at day 3 or 7. TUNEL staining was used to evaluate photoreceptor cell death. Photoreceptor cell loss was evaluated by measuring the thickness of outer nuclear layer (ONL). Microglia in the ONL were quantified in a retinal whole mount with Iba-1 antibody. RPE degeneration was also assessed in a RPE whole mount, with ZO-1 antibody.

**Results:** NaIO3 resulted in significant cell death of ARPE-19 cells. Treatment with Nec-1 resulted in better protection than treatment with Z-VAD (P<0.01). A synergistic protective effect was observed when co-treating the cells with Nec-1 and Z-VAD. Nec-1 treatment also decreased the ARPE-19 mitochondrial damage caused by NaIO3. *In vivo* administration of NaIO3 resulted in significant RPE and photoreceptor destruction with substantial inflammatory cell infiltration. RIP3 knockout animals displayed considerably less RPE and photoreceptor cell loss, as well as drastically less inflammation.

**Conclusions:** Programmed necrosis is an important cell death pathway mediating NaIO3 RPE and photoreceptor cell toxicity. Blocking the necroptosis pathway may serve as a novel therapeutic strategy for various RPE degenerative diseases.

## INTRODUCTION

Degeneration and death of the retinal pigment epithelium (RPE) and the photoreceptors are the hallmark of several vision-impairing diseases, such as age-related macular degeneration (AMD) and retinitis pigmentosa (RP) (1,2). Extensive research about these diseases has been done using a large number of both inherited and induced animal models (3–5). Sodium iodate (NaIO_3_) typically induces both RPE cell and photoreceptor death, and has been used as a model of death for both cell types (6–8). Sodium iodate initially weakens the bond between the RPE and Bruch’s membrane and leads to RPE cell and photoreceptor death It causes progressive structural damage to the retina, with swelling, thinning, disintegration of individual retinal layers, and clustering of highly reflective cellular debris (9–11). The outer retina becomes significantly disorganized and thinned. Concurrently, there is Müller cell proliferation and macrophage migration within the retina (12). On histopathology, the retina shows a mosaic pattern, with some areas relatively normal and others lacking RPE and photoreceptor cells(8). Recent studies reported that NaIO3-induced visual dysfunction is associated with oxidative stress and rapid suppression of phototransduction genes in photoreceptors, suggesting that NaIO3 can directly alter photoreceptor function and survival (6). There are several known mechanisms for the effect of NaIO_3_ on RPE cells. NaIO_3_ has been shown to react with melanin, converting the substance into a toxic compound (13). As a result, various enzymatic activities such as glycolysis, Krebs cycle, and pentose cycle are inhibited (14), while the adhesions between the sensory retina and the RPE are weakened, leading to disruption of the blood retinal barrier (15). However, the precise mechanism through which NaIO_3_ provokes photoreceptor and RPE cell death remains unclear.

Apoptosis and necrosis are two major distinct pathway of cell death, both defined by morphological criteria (16,17). Apoptosis is the process of programmed cell death characterized by morphological cellular changes, including cell shrinkage, nuclear fragmentation, chromatin condensation, and chromosomal DNA fragmentation (18,19). The caspase family of proteases plays a central role in the induction phase of this process. Z-VAD is a pan-caspase inhibitor and has been widely used to study the apoptotic pathway. Necroptosis is another form of cell death that is characterized by cell swelling, rapid permeabilization of the cell membrane, release of cell contents and exposure to damage-associated molecular patterns. Traditionally, it has been described as uncontrolled and accidental necrosis. However, recent evidence shows that multiple mediators regulate this type of cell death. Two members of the receptor-interacting protein (RIP) kinase family proteins, RIP1 and RIP3, have been identified as critical mediators of necroptosis (20–23). RIP1 is a multifunctional death-domain adaptor protein that mediates both apoptosis and necrosis. RIP1 induces apoptosis when recruited to the protein complex containing Fas-associated death domain and caspase-8 (19,24,25). When caspases are either inhibited or not activated, RIP1 binds to RIP3, forming a pronecrotic complex, which is stabilized by phosphorylation of their serine/threonine kinase domains (26). This is confirmed by the finding that TNF-alpha mediated necrosis can be inhibited by a specific inhibitor of RIP1 kinase, necrostatin 1 (Nec-1). (27–29). RIP kinase-dependent necrosis has been implicated in various forms of developmental and pathological cell death, including photoreceptor cell death ((30–34). To understand the effect of blocking the necroptosis pathway on RPE and photoreceptor damage induced by NaIO_3,_ we investigated the effect of blocking RIP1 and RIP3 kinase on NaIO_3_-induced RPE and photoreceptor cell death in vitro and in vivo.

## Materials and Methods

### ARPE-19 Cell Culture and Cell Viability Assays

A human RPE cell line (ARPE-19, obtained from ATCC, Manassas, VA) was used in this study. The cells were grown in a humidified 10% CO2 atmosphere at 37◦C in Dulbecco’s MEM/F-12 (1:1) medium (Life Technologies, Grand Island, NY) containing 10% heat inactivated fetal bovine serum (Life Technologies), 100 units/mL penicillin (Life Technologies) and 100ug/mL streptomycin (Life Technologies). Cells were pre-treated with Necrostatin-1 (Sigma-Aldrich) or z-Vad (Sigma-Aldrich) at 30 μM for 1 hour then co-treated with different concentrations of NaIO_3_ for various durations to examine the potential mechanisms of cell death and protection. Cell viability was examined using 3-(4,5-dimethylthiazol-2-yl)-2,5-diphenyltetracolium bromide (MTT) (Life Technologies, Eugene, OR) and lactate dehydrogenase (LDH) (CELL BIOLABS, INC) assay. After each treatment, the culture medium in each plate was replaced with MTT (0.5mg/mL) dissolved in 100uL of PBS per well and incubated for an additional 3 hours. The formed formazan crystals were then dissolved by the addition of 0.1 mL isopropanol (Acros Organics, Fair Lawn, NJ) with 0.04 N HCl (Sigma-Aldrich) to each well. Absorbance at 590nm was measured using the SpectraMax 190 Microplate Reader (Molecular Devices, Sunnyvale, CA). LDH release was measured by using an LDH assay kit (Cayman Chemical, Ann Arbor, MI) as per manufacturer’s instructions.

### Mitochondrial Detection

Cells were grown on cover slips inside a Petri dish, and filled with the appropriate culture medium. When the cells reached 80% confluence, the medium was carefully removed. The cells were stained with MitoID Red Detection Kit as per protocol (Enzo Life Science). Stained cells were analyzed by wide-field confocal microscopy (60X magnification), using a standard Rhodamine or Texas Red filter set to visualize the mitochondria. The nuclei were stained with DAPI.

### Animals

All animal experiments followed the guidelines of the ARVO Statement for the Use of Animals in Ophthalmic and Vision Research and were approved by the Animal Care Committee of The Massachusetts Eye and Ear Infirmary. WT C57BL/6 mice were purchased from The Jackson Laboratories. Rip3−/− mice were provided by Vishva M. Dixit (Genentech, San Francisco, CA) and backcrossed to C57BL/6 mice (35). Anesthesia was achieved by intraperitoneal injection of 50 mg/kg Ketamine hydrochloride (Phoenix Pharmaceutical, Inc., St. Joseph, MO) and 10 mg/kg Xylazine (Phoenix Pharmaceutical, Inc.).

### TUNEL staining

TUNEL procedure and quantification of TUNEL- positive cells were performed using an ApopTag Fluorescein Direct in Situ Apoptosis Detection Kit (Millipore) according to the instructions of the manufacturer. For retinal cryosection immunostaining, sections through the retina (10 μm) with the optic nerve attached were pre-blocked (PBS containing 5 % donkey serum, 0.5% gelatin, 0.3% BSA, and 0.1% Triton-X). Five sections were randomly selected in each eye. The posterior pole retinas were photographed, and the number of TUNEL-positive cells in the outer nuclear layer (ONL) was counted by masked observers. The retinal area was measured by ImageJ software. The data is expressed as TUNEL-positive cells per square millimeter of retinal area.

### RPE immunohistochemistry

Under deep anesthesia, mice were perfused through the left ventricle with 10 mL of PBS followed by 10 mL of 4% paraformaldehyde. Eyes were enucleated, and the anterior segment and retina were removed under a dissecting microscope. Special attention was paid during the removal of the retina to avoid damage of the underlying RPE. Four relaxing radial incisions were made, and the remaining RPE-choroid-sclera complex was flat mounted on a glass slide. Flat mounts were air-dried for 10 minutes and incubated for 1 hour with blocking solution (5% donkey serum, 0.3% bovine serum albumin, 0.3% Triton-X). Then, primary antibody against ZO-1(1:100, Invitrogen, USA) was incubated overnight at 4°C in a moisture chamber. After three washes, retinas were incubated with a donkey antibody against rabbit IgG conjugated to Alexa Fluor 594, diluted in blocking medium (1:400; Invitrogen) overnight at 4°C and flat mounted onto slides using Vectashield mounting medium after making four radial slits. The RPE was examined by a fluorescein microscope (Zeiss Axiovsion version 4.8.2, Germany). We measured the length of intact RPE in 5 different areas per quadrant and in all 4 quadrants by Image J software (developed by Wayne Rasband, National Institutes of Health, Bethesda, MD; available at http://rsb.info.nih.gov/ij/index.html) and compared RIP3 KO mice to control mice 3 days after SI injection.

### Retinal immunohistochemistry

For whole-mount immunostaining, fixed retinas were washed several times in fresh PBS, transferred to multi-well plates, and blocked in 5 % donkey serum (Sigma-Aldrich, St. Louis, MO) with 0.5% Triton X-100 in 0.1 ml PBS overnight. The retinas were then incubated in the same medium containing rabbit anti-Iba-1 antibody (1:500; Wako Chemicals USA Inc., Richmond, VA) for 1 day at 4°C on a shaking platform. After three washes, retinas were incubated with a donkey antibody to rabbit IgG conjugated to Alexa Fluor 488 diluted in blocking medium (1:400; Invitrogen) overnight at 4°C and flat mounted onto slides using Vectashield mounting medium after making four radial slits. The neuroretinas were examined with a fluorescein microscope (Zeiss Axiovsion version 4.8.2, Germany).

### Statistical analysis

Results are expressed as mean values ± standard error of mean (SEM). Statistical analysis was performed with the non-parametric Mann-Whitney U-test (SPSS Statistics 17.0, Chicago, IL). A p-value of less than 0.05 was considered statistically significant.

## RESULTS

### RIP1 inhibitor, Necrostatin-1, and Caspase inhibition blocked RPE cell death induced by NaIO_3_

To test the toxicity of NaIO3 to ARPE-19 in vitro, cells were treated with various concentrations of NaIO3 and cell death was determined by the MTT assay. As shown in Fig1 A, Sodium Iodate applied for 4 hrs to ARPE-19 cells lead to significant cell death at concentrations over 4 mg/ml. To test the role of Caspase and/or RIPK inhibition, ARPE-19 cells were pre-incubated with 4 mg/ml of NaIO3 in the presence or absence of pan-caspase inhibitor zVAD, RIPK inhibitor Nec-1, or combination of both inhibitors. As shown in Figure 1B and 1C, both Necrostatin-1 and z-VAD decreased cell death induced by NaIO3, and the combination of both inhibitors showed the greatest effect.

**Figure 1.**
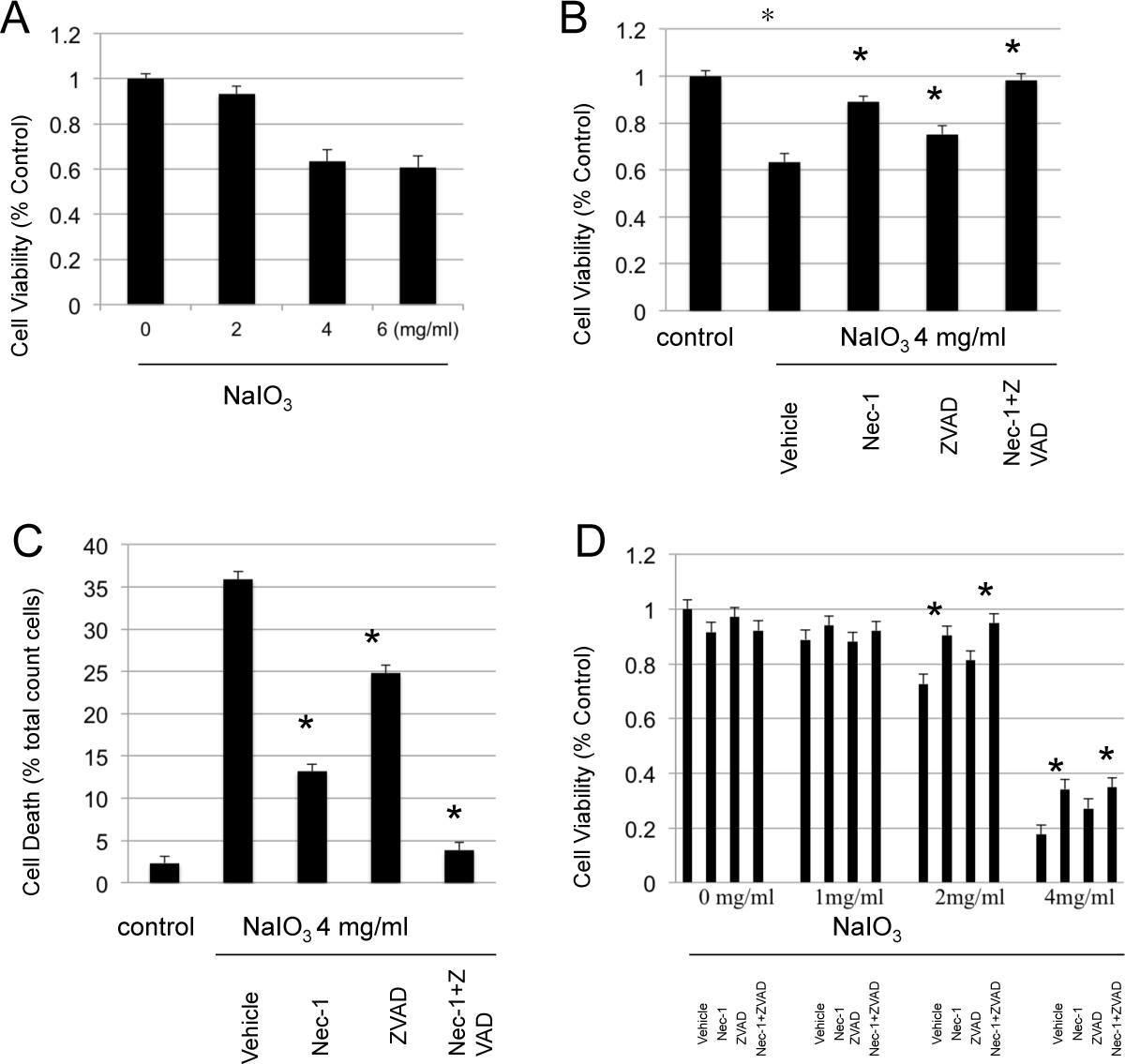
Necrostatin-1 inhibits RPE cell death induced by NaIO_3_. **A**. ARPE19 cells were treated with different concentration of NaIO3 for 4 hours. The cell death was determined with MTT assy. **B.** ARPE19 cells were treated with 4mg/ml of NaIO3 for 1 hour. The cells were treated with NaIO_3_ with/without Necrostatin-1 and z-VAD for extra 3 hours. The death was determined with MTT assy. **C.** ARPE19 cells were treated in the same way as above. The cell death were determined by LDH assay.

### Necrostatin-1 blocks mitochondrial fragmentation associated with necrosis induced by NaIO_3_

To monitor the mitochondrial morphological changes associated with NaIO_3_-induced RPE necrosis in detail, ARPE-19 cells were exposed to 4 mg/ml of NaIO_3_ for 4 hours in the presence or absence of RIP1 Kinase inhibitor Nec-1 for the last 3 hours. Mitochondria and nuclei were detected with Mito-ID Red Detection kit and DAPI staining, respectively. As seen in Figure 2, after 4 hours of NaIO3 treatment, mitochondria became more punctate and fragmented (Figure 2B). Necrostatin-1 decreased mitochondrial fragmentation induced by NaIO3, however these changes were not completely inhibited (Figure 2C).

**Figure 2.**
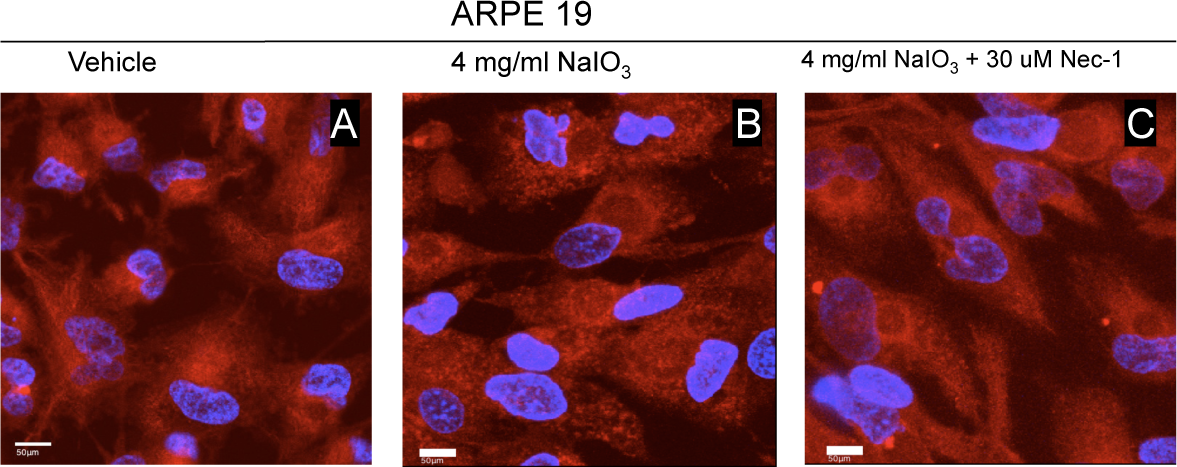
Necrostatin-1 blocks mitochondrial fragmentation associated with necrosis induced by NaIO_3_. **A**. ARPE-19 cells without *NaIO*_3_ and necrostatin-1 treatment. **B**. ARPE-19 cells were treated with 4mg/ml of NaIO_3_ for 4 hour. **C**. ARPE-19 cells were treated with 4mg/ml of NaIO_3_ for 1 hour and then continually treated with 30uM of Necrostain-1 together with NaIO3 for another 3 hours. Red represents mitochondria stained with Mito-ID Red. Blue represents nuclear with a nuclear marker DAPI.

### RIP3 deficiency diminishes RPE death after Sodium iodate administration in vivo

To further examine the role of RIP-Kinases in NaIO_3_-induced RPE toxicity, we administered NaIO_3,_ (35-mg/kg) intraperitonealy to both WT C57BL/6 mice and RIP 3 -/-mice. The retina and RPE were examined on day 3, and 7 after NaIO_3_ injection. NaIO_3_ administration resulted in marked RPE damage with a distinct demarcation line of normal RPE at the peripheral retina in both WT and RIP 3 deficient mice. On the third and seventh day after NaIO_3_ injection, we detected destruction of the RPE with preservation of the normal hexagonal pattern of ZO-1 labeling at peripheral retina, in both WT and RIP 3 -/-mice (Fig.3 E-L). When we measured the length of intact RPE on day 3 after NaIO3 injection, there was significantly increased preservation of RPE in the RIP3 -/-mice compared to the WT (618.97 ± 202.21 μm vs. 220.66 ± 48.28 μm, Mann-Whitney U test, P<0.01).

### Microglial cell activation after NaIO3

To assess inflammatory response to NaIO_3_, we performed immunofluorescence for the microglial marker Iba-1. Immunostaining with the Iba-1 antibody revealed activated microglial cells in the retina overlying the damaged RPE, while no microglia were detected beyond the demarcation line (Fig. 3 M-P, arrowhead). At day 3 after NaIO3 injection, activated amoeboid microglial cells with retracted processes and rounded cell bodies were seen in the ONL, and significantly more in the WT mice than the RIP3 -/-mice (Fig. 4 A, B, and E, Mann-Whitney U test, P< 0.01). By day 7, microglial infiltration was decreased in both WT and RIP 3-/-mice with no difference in WT mouse retinas and RIP 3-/-mouse retinas (Fig. 4 C, D, and E; Mann-Whitney U test, P =0.145). Therefore, RIP3 deficiency inhibits microglial cell activation by NaIO3 and corresponding RPE alteration.

**Figure 3.**
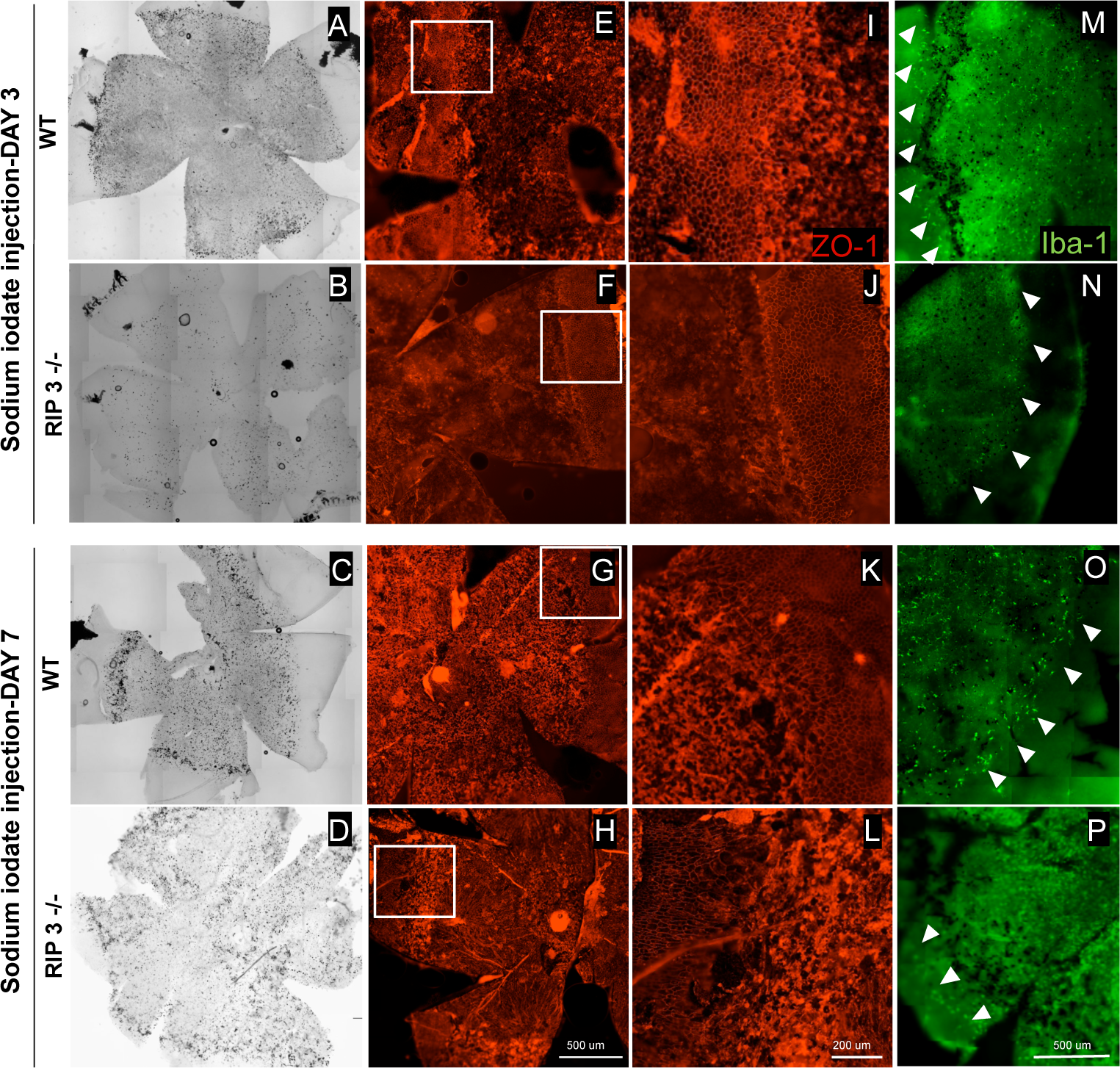
NaIO3 induced RPE/ photoreceptor cell layer damage in posterior retina with sparing in the peripheral retina. **A-D**, Macroscopic examination of retinal flat mounts from wild-type (WT) and Rip 3-/-mice showed increased RPE pigmentation in WT compared to Rip 3 deficient mice at day 3 and 7 after NaIO3 intraperitoneal injection. **E-L**, NaIO3 injection induced localized RPE destruction in posterior pole in both WT and Rip 3 -/-mice as indicated by ZO-1 staining. **M-P**, Microglial cell activation was found localized at areas of dysfunctional RPE cell membranes more in WT compared to the Rip 3 deficient retina. Scale bars: 500 μm (E-H), 200 μm (I-L), 500 μm (M-P).

**Figure 4.**
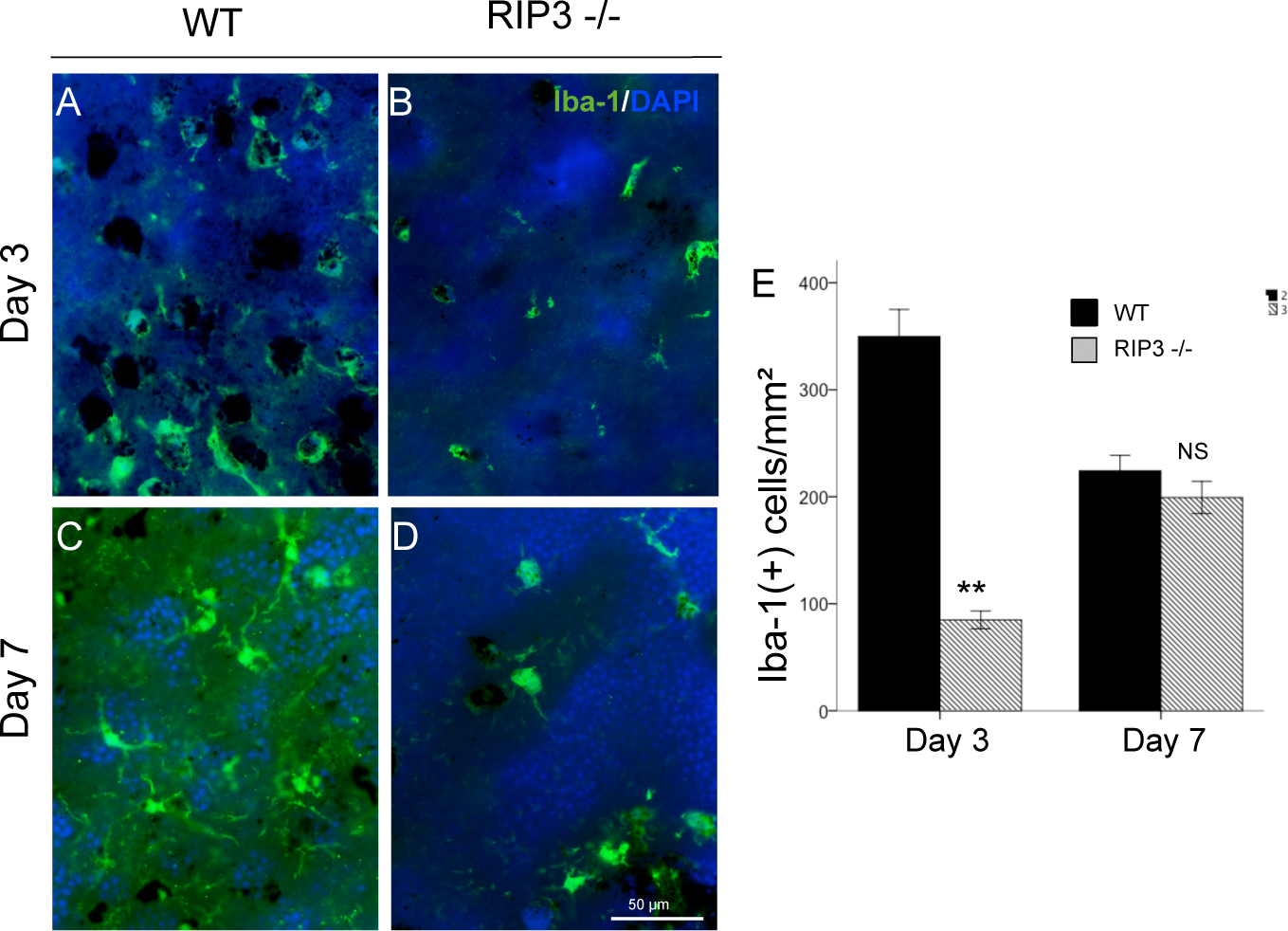
Decreased microglial cell activation after NaIO_3_ injection in RIP 3 -/-mice. **A-D**, Immunofluorescence for Iba-1 (green) and DAPI(blue) in ONL of retinal flatmount. **E**, Quantification of Iba-1+ microglia in WT and Rip3−/− mice at day 3 and 7 after NaIO_3_ injection. Microglial infiltration was significantly reduced in Rip3−/− retinas compared with WT retinas at day 3 after NaIO3 injection (A and B), whereas no difference in microglial activation was observed at day 7 (C and D). Mann-Whitney U test, **P < 0.01; NS, not significant. Data are expressed as the mean ± SEM (n = 6 for each group). Scale bar, 50 μm.

### RIP3 kinase deficiency prevents photoreceptor death caused by NaIO_3_ injection

Next, we assessed photoreceptor death after NaIO_3_ administration by TUNEL staining. By day 3, increased TUNEL-positive cells were noted in the ONL in both WT and RIP 3-/-mice, which decreased by day 7 (Fig. 5 A-D). On both day 3 and 7, the number of TUNEL-positive cells in ONL noted in the RIP 3 -/-mice was significantly reduced, as compared to the WT (Mann-Whitney U test, P <0.001 and P <0.001 at each point respectively; Fig. 5E). *Rip3* deficiency significantly prevented the reduction of the ONL thickness at 3 and 7 days after NaIO_3_ injection (P <0.001 and P < 0.001, respectively).

**Figure 5.**
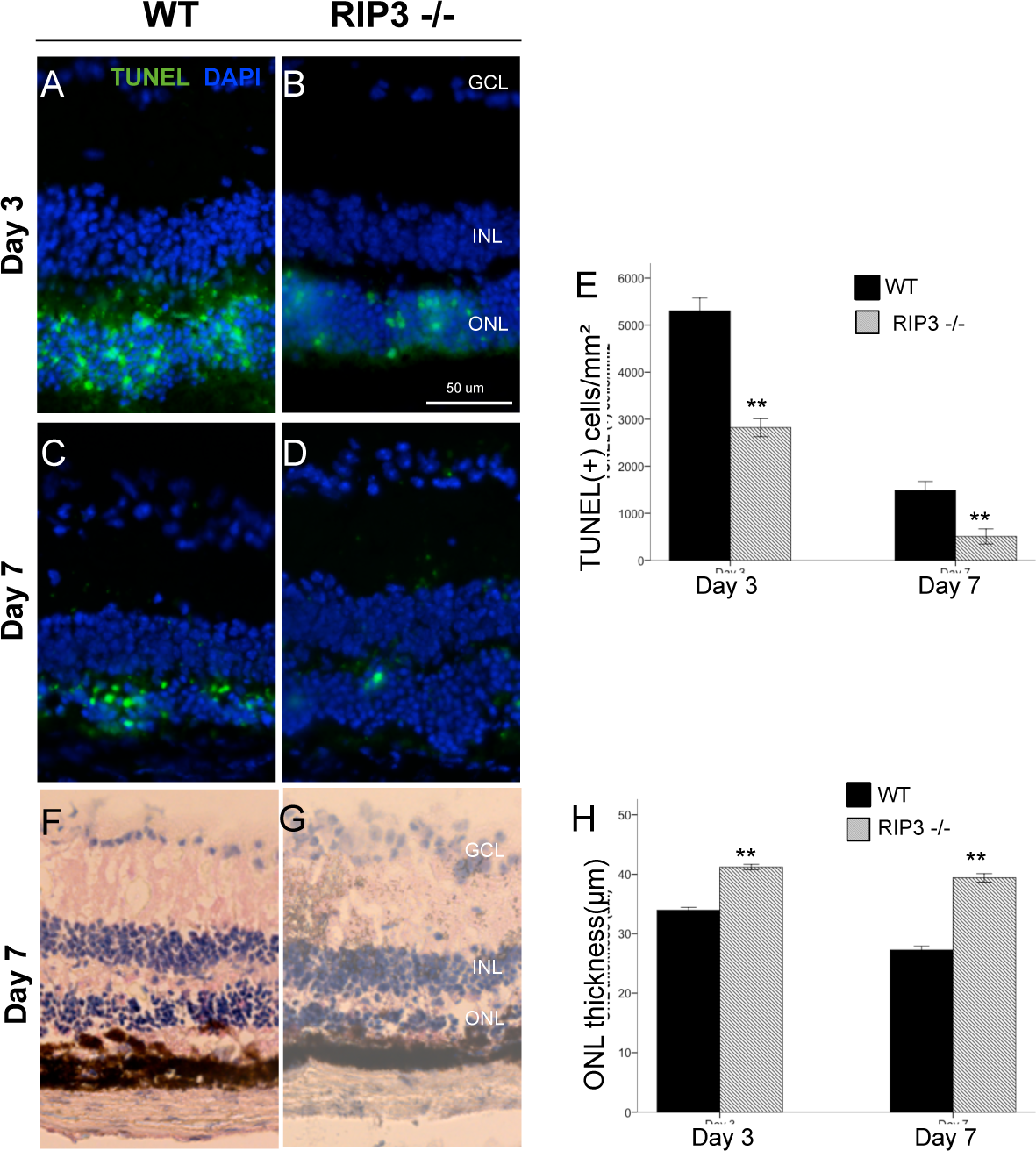
Rip 3 deficiency prevents apoptosis in photoreceptor after NaIO_3_ injection. **A-D**, TUNEL (green) and DAPI (blue) staining at day 3 and 7 after NaIO3 injection in WT mice and Rip3−/− mice. GCL, ganglion cell layer; INL, inner nuclear layer; ONL, outer nuclear layer. Quantification of TUNEL-positive photoreceptors (**E**) shows significantly decreased apoptosis in the photoreceptor layer with Rip 3 deficiency. Retinal histology (**F-G**) and ONL thickness quantification (**H**) at day 3 and 7 after NaIO_3_ injection in WT mice and Rip3−/− mice show significant preservation of ONL with Rip 3 deficient mice. Data are expressed as the mean ± SEM (n = 6 for each group). Mann-Whitney U test, **P < 0.01 comparing WT and Rip 3. Scale bar: 50 μm.

## Discussion

NaIO3 is a drug commonly used to induce retinal degeneration via oxidative stress and subsequent cell death in animal models by apoptosis (7,8,12,36,37). More specific, recent evidence suggests that early photoreceptor damage may be a key event in the pathogenesis of NaIO3-induced retinal degeneration (6). This is the first study to investigate the role of RIPK regulated necrosis in NaIO_3_-induced RPE and photoreceptor degeneration. In addition, we provide evidence that both apoptosis and necroptosis pathways are involved in RPE cell death induced by NaIO3. Our results show that treatment of human retinal pigment epithelial (ARPE-19) cells for 4 hours with concentrations of NaIO3 greater than 2 mg/mL leads to cell death, Furthermore, this effect can be blocked partially by RIP1 specific inhibitor and pan caspases inhibitor with synergistic effect shown with both Necrostatin-1 and z-VAD together. In addition, we have also found that RIP3 deficiency reduces photoreceptor cell death after NaIO_3_ administration in vivo, which was also associated with a decreased inflammatory response.

Interestingly, our results show that Nec-1 confers greater protection on ARPE-19 cells than z-VAD (Fig1). While z-VAD is a pan-caspase inhibitor, Nec-1 is a necrosis-specific inhibitor which does not affect apoptosis (30). This suggests that necroptosis pathway may be more important than apoptosis in RPE cell death caused by NaIO_3_. A number of studies have shown necrosis to be a major mechanism of RPE death during oxidative stress (30,38,39), and it is now recognized as a regulated process mediated by receptor-interacting protein (RIP) kinases (33,40). Nec-1 inhibits RIP1 kinase phosphorylation during necrosis, which is required for the formation of the RIP1-RIP3 complex (necrosome) and RIP3 kinase phosphorylation (26,28,41). Our data also support RIP kinase-mediated necrosis as a cell death pathway in ARPE-19 cells undergoing stress via NaIO3 administration. Additionally, it evidences that simultaneous inhibition of both RIP kinase and caspase pathways lead to almost complete cellular protection (Figure 1B and 1C).

Mitochondrial fragmentation by membrane permeabilization and the subsequent release of mitochondrial proteins into the cytoplasm is one of the downstream effects of both the apoptosis and necrosis pathways (42). We have shown that ARPE-19 cells treated with NaIO_3_ demonstrate significant mitochondrial fragmentation (Figure 2B), denoted by the transformation of their normally tubular morphology into small particles scattered throughout the cell. This process is partially blocked by Nec-1 (Figure 2C), suggesting that it can occur, at least in part, via the necrotic pathway.

RIP3 Kinase is a key regulator of RIP1 kinase phosphorylation and necrotic signaling. Activation and increased expression of RIP3 is a hallmark of necrosis (41,43),(30). Our histopathological findings in Figure 3 show that mice lacking RIP3 show decreased RPE cell damage and photoreceptor cell death after injection of NaIO_3_ at day 3. This suggests that necrosis is a major mechanism of cell death by NaIO_3,_ and that RIP3 is a key regulator of this process. These data supports our previous findings, in which RIP3 deficiency attenuated both photoreceptor death after retinal detachment and PolyI:C injection (30).

In contrast to apoptosis, necrosis is associated with significant inflammatory response (44). Consistent with this, it has been suggested that necrotic RPE cells can be associated with inflammatory response in neighboring cells (38). After NaIO_3_ injection, significant microglial cell infiltration was noted in the ONL. Interestingly, knockdown of *Rip3* inhibited this infiltration on day 3, but not on day 7, suggesting that it may have a regulatory function in the early stages of the inflammatory response.

In summary, our study shows that NaIO3 injury to RPE and photoreceptors in addition to apoptosis RIP-mediated programmed necrosis is another essential pathway for cell loss. Furthermore, NaIO3 induces an inflammatory response that is partially blocked by *Rip3* deletion. Therefore, blocking necroptosis pathway along with caspase mediated apoptosis, may be helpful in the management of various toxic and degenerative retina and RPE diseases.

## Acknowledgments

This study is supported by National Institutes of Health Grants EY12850 and NEI R21EY023079-01/A1 (DGV), the Yeatts Family Foundation, a 2013

Macula Society Research Grant award (DGV) and an RPB Physician Scientist Award (DGV), Loeffler Family fund and a Lions unrestricted fund to MEEI.

---

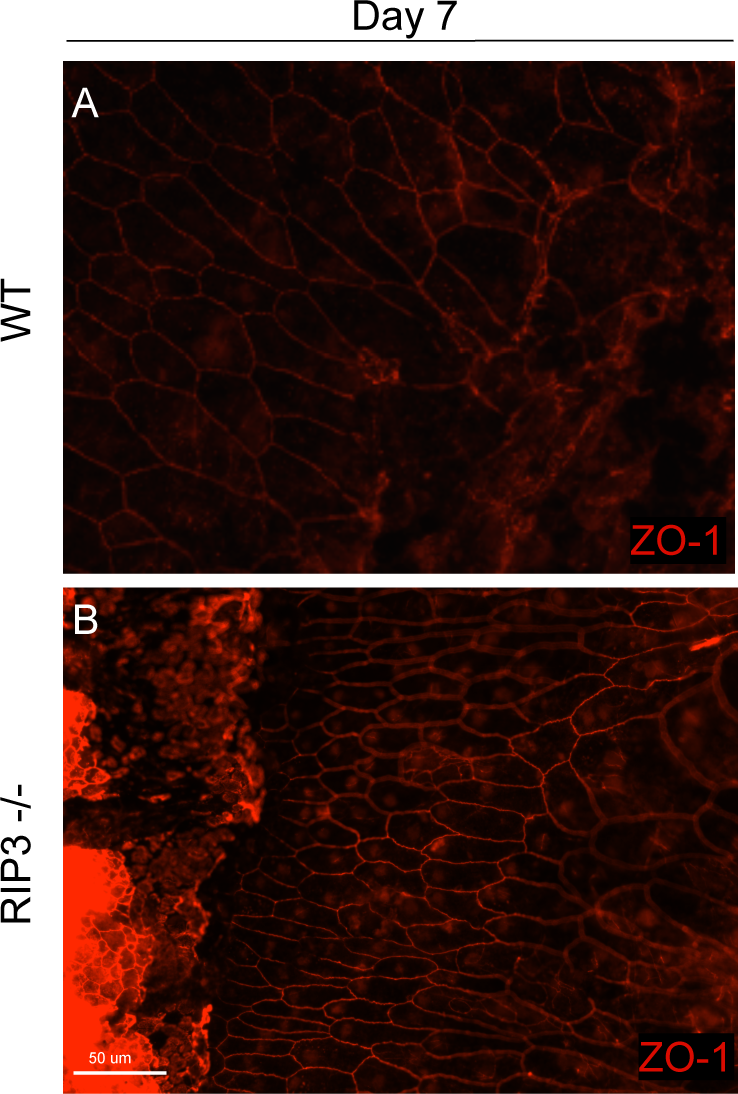

